# An immunodominance hierarchy exists in CD8^+^ T cell responses to HLA-A*02:01-restricted epitopes identified from the non-structural polyprotein 1a of SARS-CoV-2

**DOI:** 10.1101/2020.09.18.304493

**Authors:** Akira Takagi, Masanori Matsui

## Abstract

COVID-19 vaccines are being rapidly developed and human trials are underway. Almost all of these vaccines have been designed to induce antibodies targeting spike protein of SARS-CoV-2 in expectation of neutralizing activities. However, non-neutralizing antibodies are at risk of causing antibody-dependent enhancement. Further, the longevity of SARS-CoV-2-specific antibodies is very short. Therefore, in addition to antibody-induced vaccines, novel vaccines on the basis of SARS-CoV-2-specific cytotoxic T lymphocytes (CTLs) should be considered in the vaccine development. Here, we attempted to identify HLA-A*02:01-restricted CTL epitopes derived from the non-structural polyprotein 1a of SARS-CoV-2. Eighty-two peptides were firstly predicted as epitope candidates on bioinformatics. Fifty-four in 82 peptides showed high or medium binding affinities to HLA-A*02:01. HLA-A*02:01 transgenic mice were then immunized with each of the 54 peptides encapsulated into liposomes. The intracellular cytokine staining assay revealed that 18 out of 54 peptides were CTL epitopes because of the induction of IFN-γ-producing CD8^+^ T cells. In the 18 peptides, 10 peptides were chosen for the following analyses because of their high responses. To identify dominant CTL epitopes, mice were immunized with liposomes containing the mixture of the 10 peptides. Some peptides were shown to be statistically predominant over the other peptides. Surprisingly, all mice immunized with the liposomal 10 peptide mixture did not show the same reaction pattern to the 10 peptides. There were three pattern types that varied sequentially, suggesting the existence of an immunodominance hierarchy, which may provide us more variations in the epitope selection for designing CTL-based COVID-19 vaccines.

**Importance:** For the development of vaccines based on SARS-CoV-2-specific cytotoxic T lymphocytes (CTLs), we attempted to identify HLA-A*02:01-restricted CTL epitopes derived from the non-structural polyprotein 1a of SARS-CoV-2. Out of 82 peptides predicted on bioinformatics, 54 peptides showed good binding affinities to HLA-A*02:01. Using HLA-A*02:01 transgenic mice, 18 in 54 peptides were found to be CTL epitopes in the intracellular cytokine staining assay. Out of 18 peptides, 10 peptides were chosen for the following analyses because of their high responses. To identify dominant epitopes, mice were immunized with liposomes containing the mixture of the 10 peptides. Some peptides were shown to be statistically predominant. Surprisingly, all immunized mice did not show the same reaction pattern to the 10 peptides. There were three pattern types that varied sequentially, suggesting the existence of an immunodominance hierarchy, which may provide us more variations in the epitope selection for designing CTL-based COVID-19 vaccines.

## Introduction

In December 2019, the coronavirus disease 2019 (COVID-19) caused by the severe acute respiratory syndrome coronavirus 2 (SARS-CoV-2) was firstly identified in Wuhan, Hubei province, China. Since then, its subsequent spread of global infection has still continued to gain momentum. As of September 16th, 2020, the COVID-19 pandemic has infected more than 29.4 million people around the world and caused more than 931,000 deaths. Although the clinical symptom is varied from asymptomatic or mild self-limited infection to severe life-threating respiratory disease, the mechanism of disease outcome remains unclear. Many nations are struggling to find appropriate preventive and control strategies. However, there are no vaccines or antiviral drugs available for the treatment of this infectious disease.

There are seven coronaviruses that infect humans. In addition to SARS-CoV-2, SARS-CoV and middle-east respiratory syndrome coronavirus (MERS-CoV) cause severe pneumonia, whereas the other four human coronaviruses including HCoV-229E, -NL63, -OC43 and -HKU1 cause common cold (1). Like other coronaviruses, SARS-CoV-2 possesses a large single-stranded positive sense RNA genome (2). As shown in Fig. 1, the 5’-terminal two-thirds of the genome are composed of ORF1a and ORF1b. ORF1a encodes the polyprotein 1a (pp1a) containing non-structural proteins (nsp1-11) (Fig. 1). The remaining one-third of the genome encodes the structural proteins involving spike (S), envelope (E), membrane (M), and nucleocapsid (N) as well as accessory proteins (Fig. 1). Coronaviruses depend on S protein for binding to host cells. The host cell receptor for SARS-CoV-2 is the angiotensin-converting enzyme 2 (ACE2) which is also the receptor for SARS-CoV (3, 4).

**Fig. 1.**
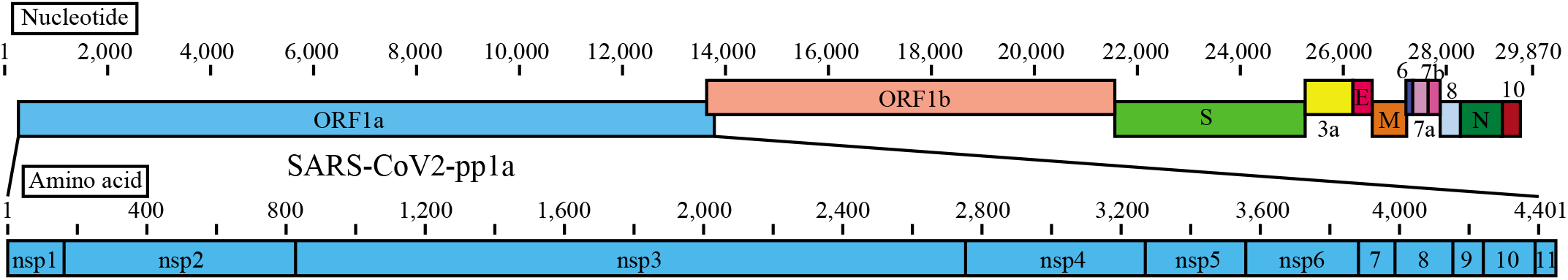
The linear diagrams of the SARS-CoV-2 genome and the protein subunits of ORF1a. The SARS-CoV-2 genome consists of ORF1a, ORF1b, Spike (S), ORF3a, Envelope (E), Membrane (M), ORF6, ORF7a, ORF7b, ORF8, Nucleocapsid (N), and ORF10. The ORF1a polyprotein is composed of eleven non-structural proteins, nsp1-nsp11.

It was shown that the increased cell numbers of antibody-secreting cells, follicular helper T cells, activated CD4^+^ T cells and CD8^+^ T cells were observed in a non-severe COVID-19 patient (5), indicating that robust immune responses can be elicited to the newly emerged, SARS-CoV-2. Therefore, the induction of protective immunity against SARS-CoV-2 is considered to help control COVID-19. Vaccines are being rapidly developed in the world, and human trials are underway for several vaccine candidates, ranging from traditional vaccines comprised of inactivated SARS-CoV-2 preparations (6) to innovative vaccines such as subunit (7), RNA (8), DNA (9), and adenoviral (10) vaccines. Almost all of these vaccines have been designed to induce antibodies targeting S protein because some antibodies specific for the receptor binding domain (RBD) of S protein may have neutralizing activities and can interfere with the binding between ACE2 on host cells and virus. In fact, it was reported that DNA vaccine encoding S protein induced neutralizing antibody in rhesus macaques and protected them from the challenge with SARS-CoV-2 (9). However, there are two major issues concerning this vaccine. One issue is that non-neutralizing and sub-neutralizing antibodies to S protein induced by this vaccine are at risk of causing antibody-dependent enhancement (ADE) (11). ADE is the phenomenon in which binding of suboptimal antibodies to viruses enhances viral entry mediated by Fc receptors into immune cells, and promotes inflammatory and tissue injury (12). ADE has been reported in the evaluation of vaccine candidates directed to S protein for SARS-CoV (13–16). Because of their similarities, these findings enable us to foresee the high risks of ADE in SARS-CoV-2 vaccines directed to S protein. It is worth noting that RBD-specific antibodies with potent neutralizing activity are extremely rare among S-specific antibodies in COVID-19-covalescent individuals (17, 18), suggesting that the development of effective vaccines might requires novel strategies to selectively target RBD of SARS-CoV-2. On the other hand, Passive immunization of RBD-specific monoclonal antibodies obtained from convalescent individuals might be safe and effective for the elimination of SARS-CoV-2 (19–21), but much more expensive to produce for worldwide use than vaccines (22). The other issue is the short longevity of neutralizing antibodies to SARS-CoV-2. It was previously reported that the SARS-CoV-specific antibodies are short-lived for at most about 2-3 years (23, 24) in comparison with SARS-CoV-specific memory T cells (25). The longevity of SARS-CoV-2-specific antibodies are likely to be even shorter, as indicated by antibody titers being undetectable or approaching baseline in the majority of SARS-CoV-2-infected individuals after 2-3 months post onset of symptoms (26). Taken together, these data strongly suggest that it does not seem right to rely too much on just the S-specific antibody-induced vaccine to control the COVID-19 pandemic.

In the viral infection, CD8^+^ cytotoxic T lymphocytes (CTLs) play an important role for the clearance of virus as well as neutralizing antibodies. CTLs recognize virus-derived peptides in association with major histocompatibility complex class I (MHC-I) molecules on the surface of antigen presenting cells and kill virus-infected target cells. It was reported that CD4^+^ T cells and CD8^+^ T cells are decreased in proportion to the disease severity and are exhausted in severe COVID-19 patients (27, 28), suggesting the significance of CD8^+^ CTLs in the clearance of SARS-CoV-2. Furthermore, SARS-CoV-specific memory T cells persisted up to 11 years (25), predicting the long life of SARS-CoV-2-specific memory T cells. Therefore, in addition to antibody-induced vaccines, novel vaccines based on SARS-CoV-2-specific CTLs should also be considered in the future vaccine development for the prevention and disease control of COVID-19.

For the development of CTL-based COVID-19 vaccine, we here attempted to identify HLA-A*02:01-restricted, dominant CTL epitopes derived from pp1a of SARS-CoV-2. Pp1a is a largest protein composed of 4,401 amino acids among SARS-CoV-2 proteins, and therefore, it seems highly possible to find dominant epitopes in this protein. Furthermore, pp1a is produced first in SARS-CoV-2-infected cells, suggesting pp1a-specific CTLs could kill infected cells before the formation of mature virions. In addition, this protein is composed of 11 non-structural regulatory proteins that are highly conserved among many different coronaviruses (29). To identify CTL epitopes, we used computational algorithms, HLA-A*02:01 transgenic mice and the peptide-encapsulated liposomes. In a similar way, we previously identified CTL epitopes of SARS-CoV pp1a (30).

## Results

### Prediction of HLA-A*02:01-restricted CTL epitopes derived from SARS-CoV-2 pp1a

We firstly attempted to predict HLA-A*02:01-restricted CTL epitopes derived from SARS-CoV-2 pp1a using four computer-based programs, SYFPEITHI (31), nHLAPred (32), ProPred-I (33), and IEDB (34). Eighty-two epitopes with high scores for all four programs were selected and synthesized into 9-mer peptides (Table 1). These peptides were then evaluated for their binding affinities to HLA-A*02:01 molecule using TAP2-deficient RMA-S-HHD cells. As the half-maximal binding level (BL_50_) values of the influenza A virus matrix protein 1-derived peptide, FMP_58-66_ (35) as a high binder control and the HIV reverse transcriptase-derived peptide, HIV-pol_476-484_ (36) as a medium binder control were 2.3 μM and 80.6 μM, respectively, we defined a high binder with a BL_50_ value below 10 μM, a medium binder with a BL_50_ value ranging from 10 to 100 μM, and a low binder with a BL_50_ value above 100 μM. As shown in Table 1, 20 peptides were high binders, whereas 34 peptides were medium binders, suggesting that the bioinformatics prediction was mostly successful. In contrast, the remaining 28 peptides displayed low affinities or no binding to HLA-A*02:01 molecules (Table 1). In the following experiments, 54 peptides involving high binders and medium binders were chosen to further investigate their abilities of peptide-specific CTL induction.

**Table 1.**
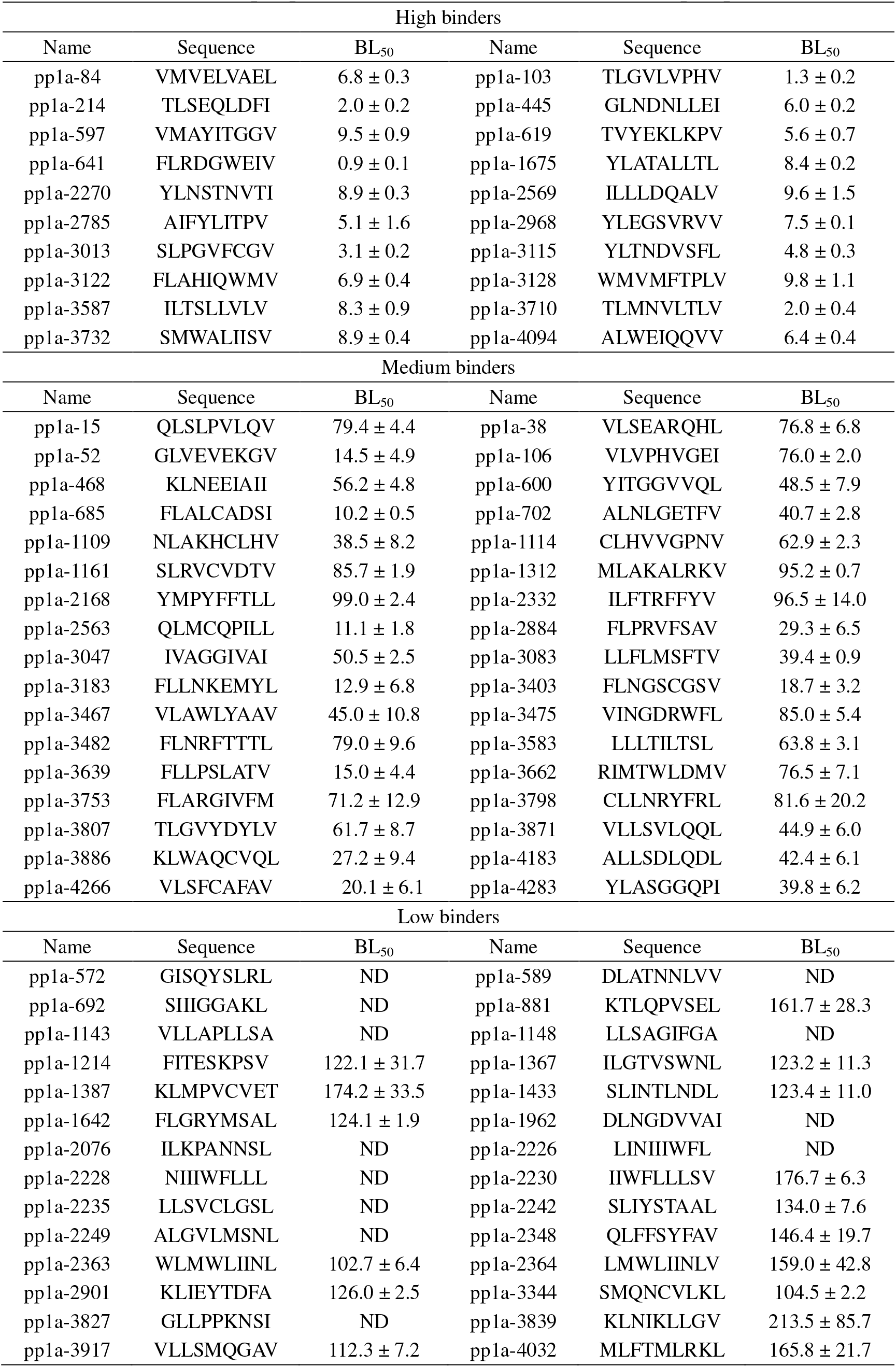
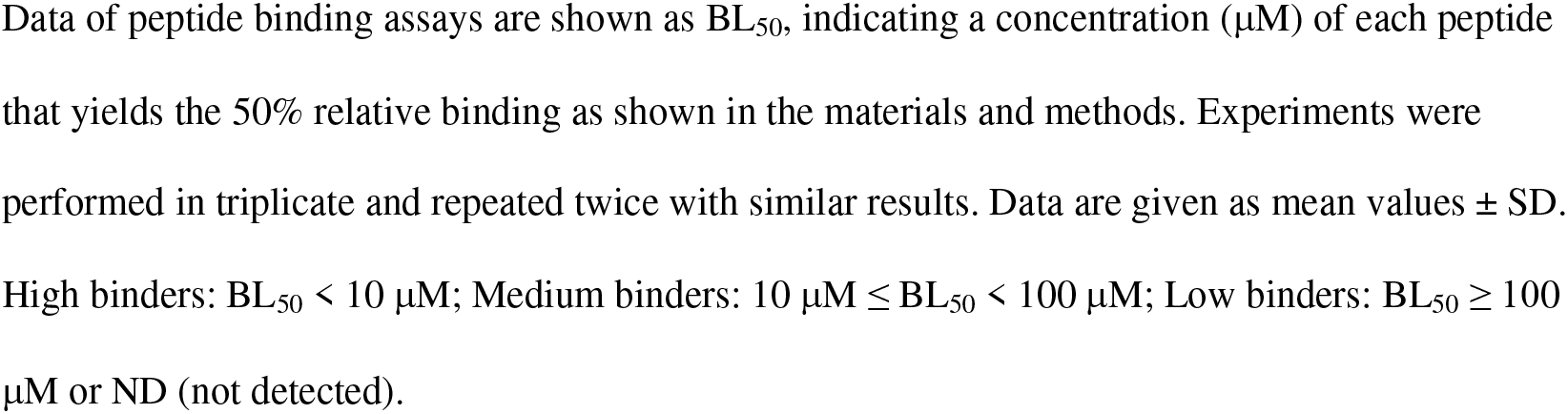
Predicted CTL epitopes for the SARS-CoV-2 non-structural polyprotein 1a

### Detection of SARS-CoV-2 pp1a-specific CD8^+^ T cell responses in mice immunized with peptide-encapsulated liposomes

Each of 54 peptides selected were encapsulated into liposomes as described in the materials and methods. HLA-A*02:01 transgenic (HHD) mice (37) were then subcutaneously (s.c.) immunized twice at a one-week interval with each of peptide-encapsulated liposomes together with CpG adjuvant. One week later, spleen cells of immunized mice were prepared, stimulated *in vitro* with a relevant peptide for 5 hours, and stained for their expression of cell-surface CD8 and intracellular interferon-gamma (IFN-γ). As shown in Fig. 2, the intracellular cytokine staining (ICS) assay showed that significant numbers of IFN-γ-producing CD8^+^ T cells were elicited in mice immunized with 18 liposomal peptides including pp1a-38, −52, −84, −103, −445, −597, −641, −1675, −2785, −2884, −3083, −3403, −3467, −3583, −3662, −3710, −3732, and −3886, revealing that these 18 peptides are HLA-A*02:01-restricted CTL epitopes derived from SARS-CoV-2 pp1a. As indicated in Table 2, multiple epitopes are located in small proteins such as nsp1 (180 aa) and nsp6 (290 aa), whereas only one epitope is seen in the large nsp3 composed of 1945 amino acids. On the other hand, the remaining 36 peptides out of 54 peptides in liposomes were not able to stimulate peptide-specific CTLs in mice (data not shown), demonstrating the necessity to generate data through wet-lab experiments. Interestingly, four epitopes including pp1a-103, −2884, −3403, and −3467 are located in the amino acid sequence of SARS-CoV pp1a as well (Table 2). pp1a-3467 was previously reported by us in the identification of SARS-CoV-derived CTL epitopes (30). However, any of 18 epitopes are not found in the amino acid sequence of either MERS-CoV or the four common cold human coronaviruses involving HCoV-OC43, HCoV-229E, HCoV-NL63, and HCoV-HKU1.

**Fig. 2.**
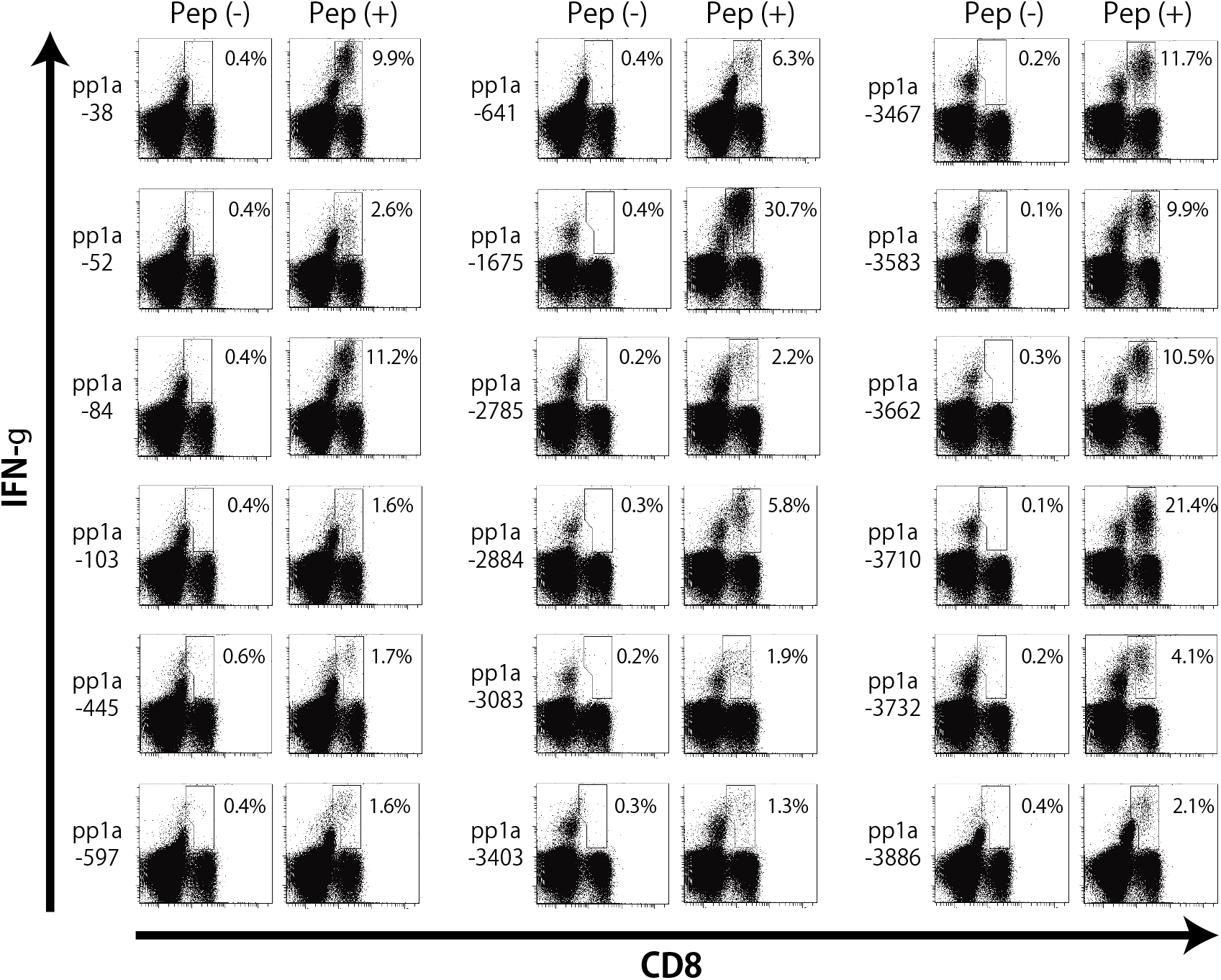
Intracellular IFN-γ staining of CD8^+^ T cells specific for peptides derived from SARS-CoV-2 pp1a. HHD mice were immunized twice with each of predicted peptides of SARS-CoV-2 pp1a in liposomes together with CpG. After one week, spleen cells were prepared and stimulated with (+) or without (-) a relevant peptide for 5 hours. Cells were then stained for their surface expression of CD8 (x axis) and their intracellular expression of IFN-γ (y axis). The numbers shown indicate the percentages of intracellular IFN-γ^+^ cells within CD8^+^ T cells. Three to five mice per group were used in each experiment, and the spleen cells of the mice per group were pooled.

**Table 2.**
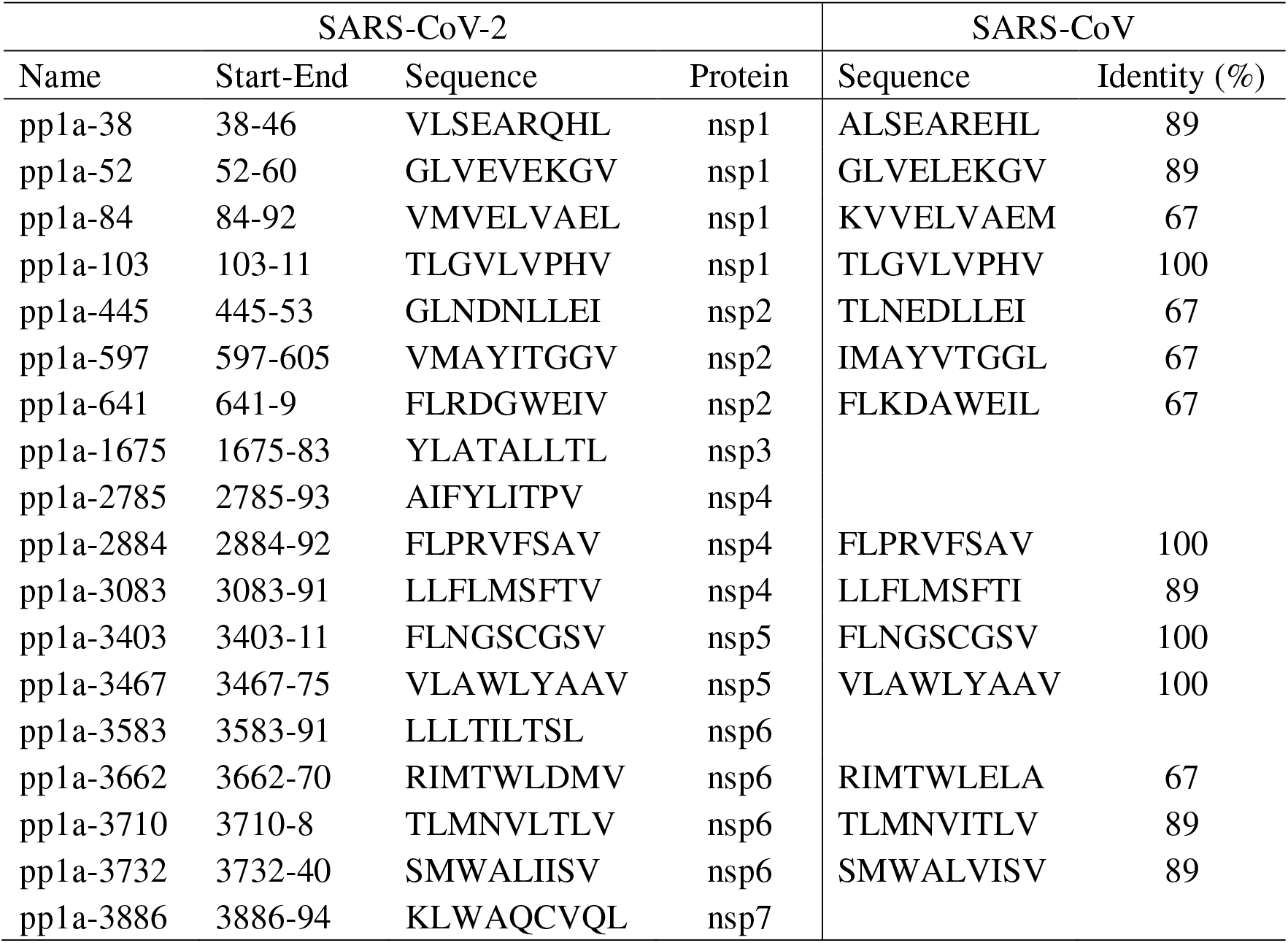
Comparison of amino acid sequences of SARS-CoV-2 pp1a CTL epitopes with the amino acid sequence of SARS-CoV

In the 18 positive peptides, 10 peptides including pp1a-38, −84, −641, −1675, −2884, −3467, −3583, −3662, −3710, and −3732 were selected for the following analyses because of the high ratios of IFN-γ^+^ cells in CD8^+^ T cells (Fig. 2).

### Identification of dominant CTL epitopes

To confirm that the 10 peptides are effective epitopes for peptide-specific CTL responses, we examined whether peptide-specific killing activities were elicited in mice with each of 10 peptides in liposomes. HHD mice were immunized s.c. twice with each of peptide-encapsulated liposomes and CpG adjuvant. One week later, equal numbers of peptide-pulsed and -unpulsed target cells were transferred into immunized mice via i.v. injection, and peptide-specific lysis was analyzed by flow cytometry. In support of the data of ICS (Fig. 2), peptide-specific killing was observed in mice immunized with any of 10 liposomal peptides (Fig. 3A).

**Fig. 3.**
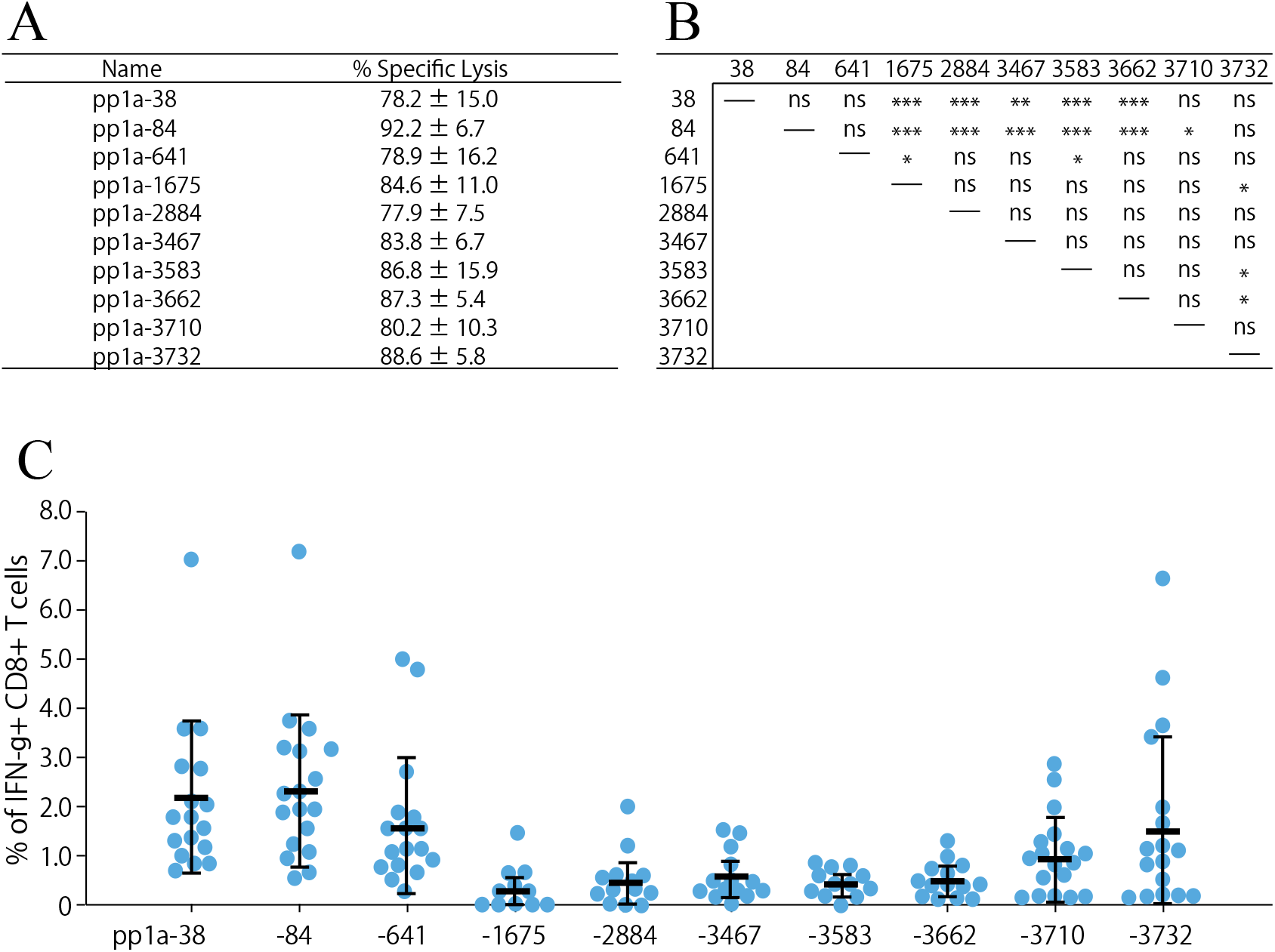
(A) *In vivo* killing activities specific for the 10 peptides. HHD mice were immunized twice with either each of the 10 liposomal peptides (pp1a-38, −84, −641, −1675, −2884, −3467, −3583, −3662, 3710, and −3732) or liposomes alone together with CpG. One week later, *in vivo* peptide-specific killing activities were measured. Three to five mice per group were used, and the data of % specific lysis are shown as the mean ± SD. (B & C) Comparison of the 10 peptides in the induction of IFN-γ^+^ CD8^+^ T cells. Seventeen HHD mice were immunized once with the mixture of 10 peptides involving pp1a-38, −84, −641, −1675, −2884, −3467, −3583, −3662, −3710, and −3732 in liposomes with CpG. After one week, spleen cells were stimulated with or without each of the 10 peptides, and intracellular IFN-γ in CD8+ T cells was stained. (C) Y-axis indicates the relative percentages of IFN-γ^+^ cells in CD8^+^ T cells which were calculated by subtracting the % of IFN-γ^+^ cells in CD8^+^ T cells without a peptide from the % of IFN-γ^+^ cells in CD8^+^ T cells with a relevant peptide. Each blue circle represents an individual mouse. Data are shown as the mean (horizontal bars) ± SD. (B) Statistical comparisons of the relative % values of IFN-γ^+^ CD8^+^ T cells among the 10 peptides in Fig. 3C were made by one-way ANOVA followed by post-hoc tests. *, *P* < 0.05; **, *P* < 0.01; ***, *P* < 0.001; ns, not significant.

We next attempted to identify dominant CTL epitopes among the 10 CTL epitopes. The same amounts of the 10 peptide solutions at an equal concentration were mixed together and encapsulated into liposomes. Seventeen mice were immunized once with the liposomes containing the mixture of 10 peptides. One week later, spleen cells were incubated with each of the 10 peptides for 5 hours, and the ICS assay was performed. As shown in Fig. 3B & C, pp1a-38 and −84 were statistically predominant over almost all other peptides in the induction of peptide-specific IFN-γ^+^ CD8^+^ T cells. Furthermore, pp1a-641 and pp1a-3732 were significantly superior to pp1a-1675/-3583 and pp1a-1675/-3583/-3662 in the stimulation of IFN-γ-producing CD8^+^ T cells, respectively (Fig. 3B).

We also examined the peptide-specific induction of CD107a^+^ CD8^+^ T cells and CD69^+^ CD8^+^ T cells. CD107a and CD69 are markers of degranulation and early activation on CD8^+^ T lymphocytes, respectively. Nine mice were immunized once with the liposomes encapsulating the mixture of the 10 peptides. After one week, spleen cells were stimulated with each peptide, and stained for their expression of CD107a or CD69 of CD8^+^ T cells. At first glance, the graphs of CD107a (Fig. 4A) and CD69 (Fig. 4B) expression were similar to that of IFN-γ expression of CD8^+^ T cells (Fig. 3C). As shown in Figs. 4A & 4C, both pp1a-38 and pp1a-84 were superior to almost all other peptides for the CD107a expression of CD8^+^ T cells. Moreover, pp1a-641 and pp1a-3732 elicited CD107a-positive T cells better than pp1a-1675/-2884/-3467 and pp1a-1675/-2884/-3467/-3583, respectively. In the case of CD69 expression (Figs. 4B & 4D), pp1a-38 and pp1a-3732 were more dominant than pp1a-1675/-2884/-3467/-3583 and pp1a-1675/-2884/-3467/-3583/-3662, respectively. Further, pp1a-84 was superior to pp1a-1675/-2884.

**Fig. 4.**
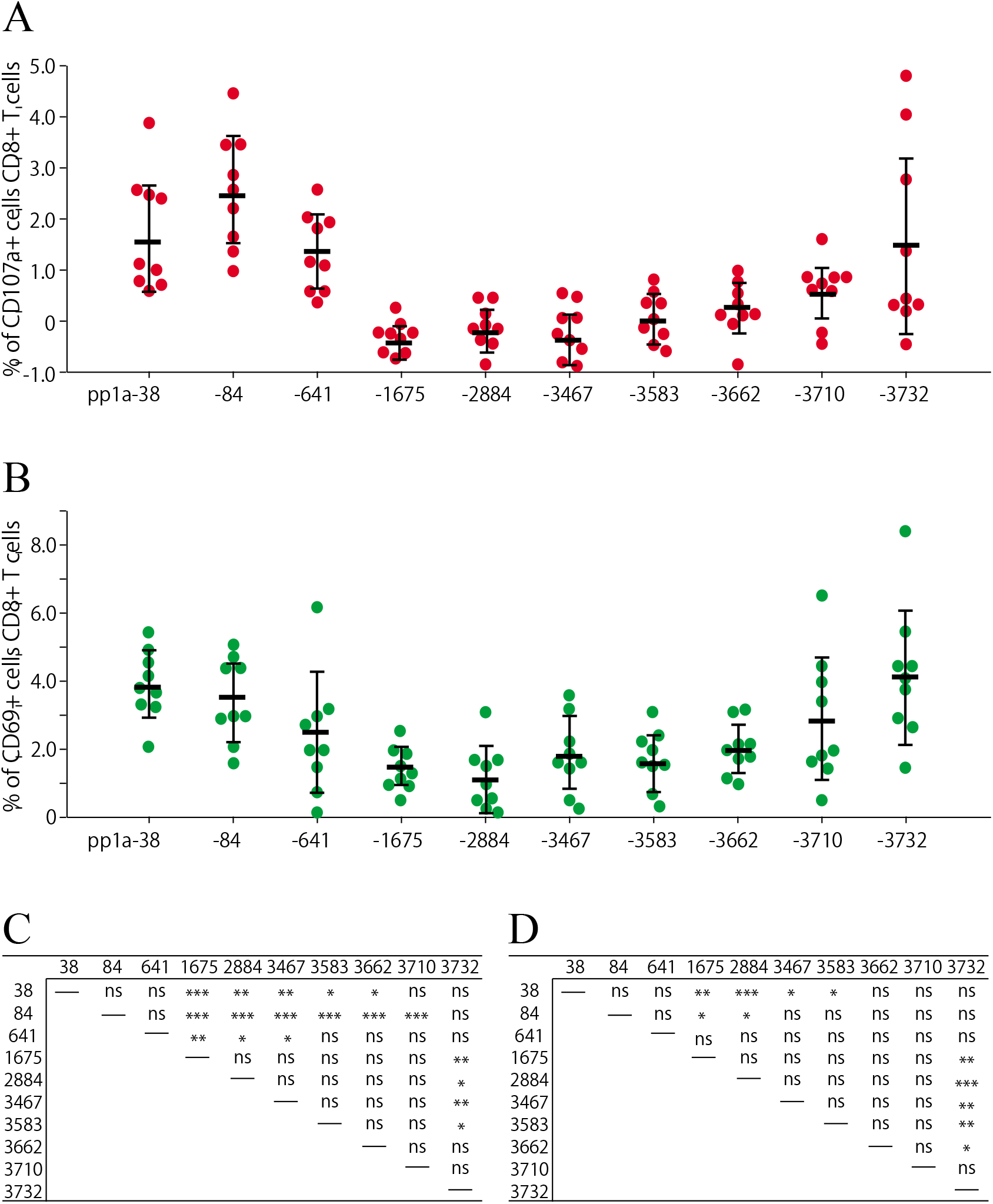
Comparison of the 10 peptides in the induction of CD107a^+^ CD8^+^ T cells (A) and CD69^+^ CD8^+^ T cells (B). Nine HHD mice were immunized once with the mixture of 10 peptides involving pp1a-38, −84, −641, −1675, −2884, −3467, −3583, −3662, −3710, and −3732 in liposomes with CpG. After one week, spleen cells were stimulated with or without each of the 10 peptides, and the expression of CD107a (A) or CD69 (B) on CD8+ T cells was analyzed. Data indicates the relative percentages of CD107a^+^ (A) and CD69^+^ (B) cells in CD8^+^ T cells which were obtained by subtracting the % of CD107a^+^ and CD69^+^ cells in CD8^+^ T cells without a peptide from the % of CD107a^+^ and CD69^+^ cells in CD8^+^ T cells with a peptide, respectively. Each red (A) or green (B) circle represents an individual mouse. Data are shown as the mean (horizontal bars) ± SD. Statistical analyses of the data among the 10 peptides in Fig. 4A and Fig. 4B were performed by one-way ANOVA followed by post-hoc tests in Fig. 4C and Fig 4D, respectively. *, *P* < 0.05; **, *P* < 0.01; ***, *P* < 0.001; ns, not significant.

Taken together, 10 peptides differed significantly in their ability to induce SARS-CoV-2 pp1a-specific CTLs when mice were immunized with the mixture of 10 peptides in liposomes. Thus, some peptides were found to be dominant CTL epitopes although each peptide alone of the 10 peptides has the capability to efficiently activate peptide-specific CTL (Figs. 2 & 3A).

### Existence of an immunodominance hierarchy

The data in Fig. 5 indicate reactivity to the 10 peptides in each of 15 mice immunized with the mixture of the 10 peptides in liposomes. Each graph represents reactivity of an individual mouse (Fig. 5). Unexpectedly, all mice did not show the same reaction pattern against the 10 peptides. It looks like there were roughly three types that varied sequentially in terms of the reaction pattern to the 10 peptides. Type A is a group of mice in which pp1a-38, −84, −641-specific IFN-γ^+^ CD8^+^ T cells were predominantly elicited, whereas the remaining 7 peptides were not able to activate peptide-specific IFN-γ^+^ CD8^+^ T cells. In the case of type B, pp1a-3732 stimulated peptide-specific IFN-γ^+^ CD8^+^ T cells as well as pp1a-38, and −84. In addition to these three peptides, several other peptides also induced IFN-γ^+^ CD8^+^ T cells in Type C. These data suggest that there seems to be an immunodominance hierarchy composed of three stages in CD8^+^ T cell responses to the 10 peptides. The immunodominance hierarchy may provide us more variations for designing CTL-based COVID-19 vaccines.

**Fig. 5.**
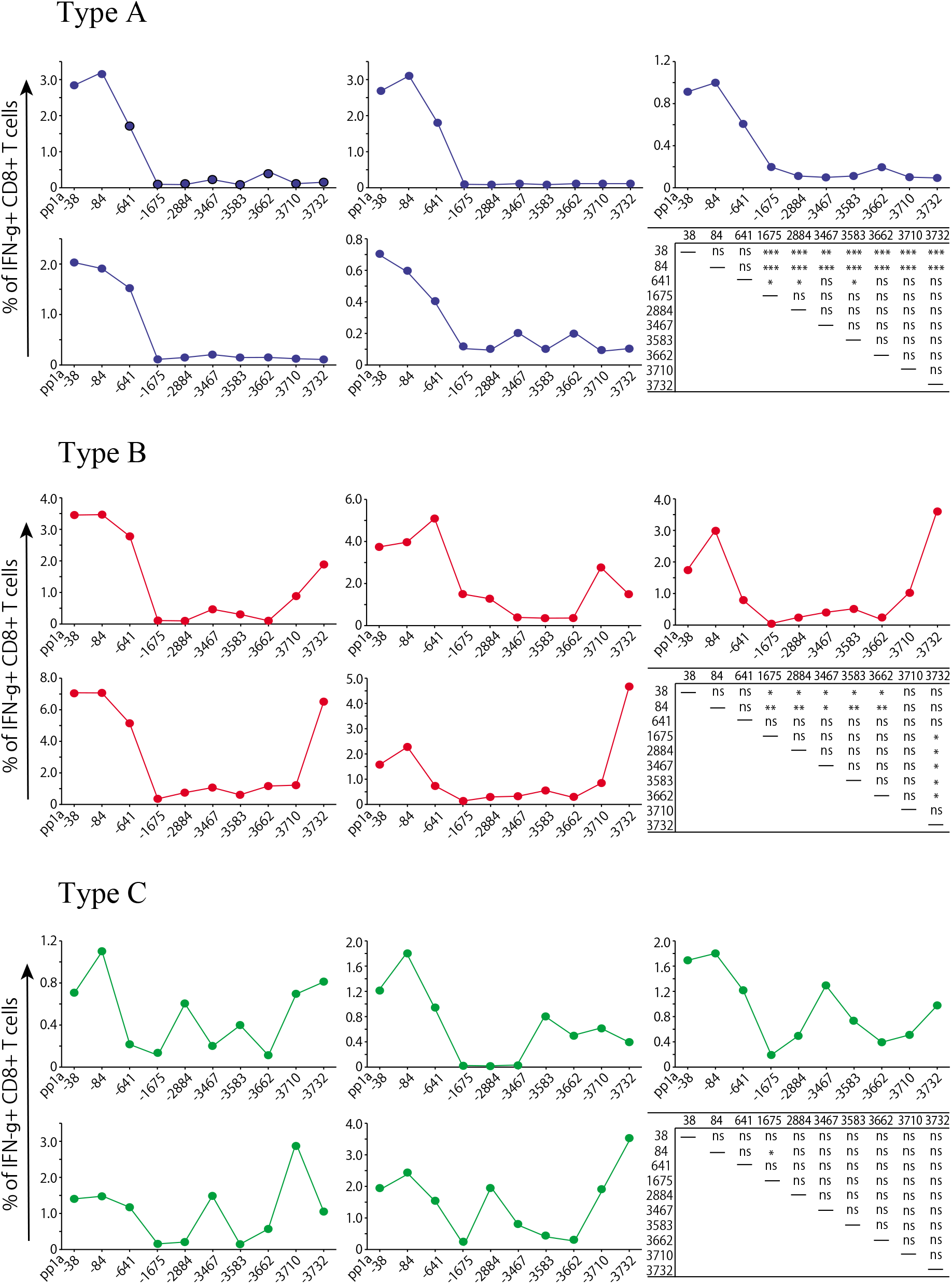
Three types of reactivity in mice immunized with the mixture of the 10 peptides. Fifteen mice were immunized once with the mixture of 10 peptides including pp1a-38, −84, −641, −1675, −2884, −3467, −3583, −3662, −3710, and −3732 in liposomes with CpG. After one week, spleen cells were stimulated with or without each of the 10 peptides, and intracellular IFN-γ in CD8+ T cells was stained. Based on the reactivity pattern to the 10 peptides, fifteen mice were divided into three types, A-C. Each graph represents reactivity of an individual mouse. Y-axis indicates the relative percentage of IFN-γ^+^ cells in CD8^+^ T cells which was calculated by subtracting the % of IFN-γ^+^ cells in CD8^+^ T cells without a peptide from the % of IFN-γ^+^ cells in CD8^+^ T cells with a relevant peptide. Statistical analyses of the relative % values to 10 peptides in each type were performed by one-way ANOVA followed by post-hoc tests. *, *P* < 0.05; **, *P* < 0.01; ***, *P* < 0.001; ns, not significant.

## Discussion

After the epidemics of SARS and MERS, scientists have not succeeded yet in developing preventive or therapeutic vaccines available for re-emergence of them. In the SARS and MERS vaccine development, the full-length S protein or its S1 subunit have frequently been used as an antigen to produce anti-RBD neutralizing antibodies. However, these vaccine candidates provided partial protection against virus challenge in animal models accompanied by safety concerns such as ADE (1). Furthermore, antibody responses to coronaviruses rapidly wane following infection or immunization (23, 24, 26). Considering the above, it should be necessary to consider CTL-based vaccine against SARS-CoV-2 to provide robust long-lived T cell memory although neutralizing antibody responses are a primary vaccine target.

In the current study, we aimed to find HLA-A*02:01-resctricted CTL dominant epitopes derived from SARS-CoV-2. Dominant epitopes induce strong immune response to eliminate a certain pathogen fast and effectively, and also contribute to make the memory T cell pool. We focused on pp1a of SARS-CoV-2 to find out CTL epitopes because pp1a is a largest and conserved polyprotein among the constituent proteins. Further, pp1a is produced earlier than structural proteins, suggesting that pp1a-specific CTLs can eliminate infected cells before the formation of mature virions. To predict CTL epitopes, we utilized bioinformatics to select 82 peptides with high scores in four kinds of computer-based programs (Table 1). In the evaluation of peptide binding, 54 peptides showed high or medium binding affinities to HLA-A*02:01 molecules, whereas the remaining 28 peptides displayed low binding affinities or no binding (Table 1). Out of them, only eighteen peptides were found to be CTL epitopes. Hence, we have to keep in mind that currently available algorithms have a limited ability to accurately predict CTL epitopes although the bioinformatics approach is very useful to quickly predict a number of epitopes in a large protein (38, 39).

Among 18 epitopes which we have identified in the current study, 4 epitopes including pp1a-103, −2884, −3403, and −3467 are present in the amino acid sequence of SARS-CoV pp1a (100% identity) (Table 2). Therefore, CTLs induced by these four epitopes could work fine for the clearance of SARS-CoV as well. In support of this data, Le Bert et al., reported that long lasting memory T cells in SARS-recovered individuals cross-reacted to N protein of SARS-CoV-2 (40). Recently, several studies found that SARS-CoV-2-reactive CD4^+^ and CD8^+^ T cells were detected in a substantial proportion of healthy donors who have never infected with SARS-CoV-2 or SARS-CoV (40–44). It is most likely that these individuals were previously infected with one of the four human coronaviruses (HCoV-229E, -NL63, -OC43 and -HKU1) that cause seasonal common cold. Nelde et al. demonstrated evidence that the amino acid sequences of several SARS-CoV-2 T cell epitopes recognized by unexposed individuals are similar to some amino acid sequences in the four seasonal common cold human coronaviruses with identities ranging from 10% to 89% (not 100% identity) (44). However, anyone has not shown evidence that people with this cross-reactivity are less susceptible to COVID-19. It may be also possible to assume that pre-existing T cell immunity might be detrimental through mechanisms such as original antigenic sin or ADE (45). In the current data, any of the 18 epitopes was not found in the amino acid sequences of the four human coronaviruses, suggesting that effective common CTL epitopes derived from pp1a, if any, might be very few.

Here, we focused on CTL epitopes restricted by HLA-A*02:01 which is the most common HLA class I allele in the world, and used highly reactive HLA-A*02:01 transgenic mice, termed HHD mice (37). Although we can use lymphocytes of SARS-CoV-2-infected individuals to identify CTL epitopes, there are mainly two advantages to using HHD mice. First, a large amount of blood of patients is required for examine many candidates of CTL epitopes, but any number of mice can be prepared for this purpose. Second, when using patients’ lymphocytes, we are only testing whether the peptide candidates are recognized by memory CTLs. When using naïve mice, however, we can find whether the epitope candidates are able to prime peptide-specific CTLs, which may be a better criterion to judge them as vaccine antigens. It is supposed that the efficient epitope for CTL recognition is not always efficient for CTL priming. However, we should take into account that the immunogenic variation in HLA class I transgenic mice may not be identical to that in humans because the antigen processing and presentation differ between them.

In the previous studies, we used peptide-linked liposomes as an immunogen (30). The surface-linked liposomal peptides were effective for peptide-specific CTL induction in mice. However, attaching peptides to the surface of liposomes followed by purifying them through the column is a fairly complicated process and time-consuming. In the current study, peptide-encapsulated liposomes were used as an immunogen. In contrast to the peptide-linked liposomes, the peptide-encapsulated liposomes are prepared by just mixing liposomes and the peptide. In addition, the peptide-encapsulated liposomes are able to prime peptide-specific CTLs in mice as efficiently as the peptide-linked liposomes. Liposome itself consisting of lipid bilayers is a very safe material for humans. Therefore, the peptide-encapsulated liposomes are considered to be promising as a CTL-based vaccine candidate.

Understanding the mechanism of immunodominance is obviously important for the development of effective vaccines. When mice were immunized with liposomes containing the mixture of 10 peptides, it was found that some peptides induced peptide-specific CTLs stronger than other peptides (Figs. 3 & 4). As shown in Figs. 3 & 4, pp1a-38 and −84 are considered to be relatively dominant in comparison with other peptides. In general, dominant epitope-specific CTLs are activated sooner and proliferate faster than subdominant epitope-specific CTLs. This immunodominance may be associated with the affinity of peptide to MHC-I molecules and the affinity of T-cell receptor (TCR) to the peptide-MHC-I complex. As shown in Table I, the peptide affinity of pp1a-84 to HLA-A*02:01 is very high (BL_50_ = 6.8 μM), while pp1a-38 is a medium binder (BL_50_ = 76.8 μM). Interestingly, the peptide affinity of pp1a-38 is lowest among the 10 peptides selected (Table I). These data suggest that the affinity of TCR to the peptide-MHC-I complex is critical for CTL immunodominance. In the selection of antigenic epitopes for the CTL-based vaccine against SARS-CoV-2, dominant epitopes such as pp1a-38 and −84 should be chosen because they produce strong CTL response to eliminate virus-infected cells effectively. However, the immunological pressure exerted by dominant epitopes may allow the epitope sequences of SARS-CoV-2 to be mutated, and therefore, a vaccine containing multiple antigenic epitopes should be recommended for a successful COVID-19 vaccine.

It was surprising that all of the genetically identical mice did not show the same reactive pattern against the 10 peptides when they were immunized with liposomes containing the mixture of 10 peptides (Fig. 5). There were roughly three pattern types, A-C, that varied sequentially, suggesting the existence of an immunodominance hierarchy composed of three stages in CD8^+^ T cell responses to the 10 peptides (Fig. 5). The differences among the three types might be explained by the timing of CTL expansion. In the type A, dominant peptides, pp1a-38, −84, and −641 presumably activated T cells more efficiently than the other peptides, and hence, dominant peptide-specific CTLs proliferate faster and curtail the expansion of CTLs specific for the other peptides. In the type B, it is considered that the expansion of dominant CTLs specific for pp1a-38, and −84 was delayed for some reason compared to that in type A, and thereby subdominant CTLs specific for pp1a-3732 could afford to expand. It is also thought that even non-dominant CTLs proliferated because the expansion of both dominant CTLs and subdominant CTLs in the type C was later than that in the type B. Although it is difficult to explain what caused difference in the timing of CTL expansion among three reaction types, it should be important to find out what it is for the development of CTL-based peptide vaccine.

In summary, we have identified 18 kinds of HLA-A*02:01-restricted CTL epitopes derived from pp1a of SARS-CoV-2 using computational algorithms, HLA-A*02:01 transgenic mice and the peptide-encapsulated liposomes. Out of 18 epitopes, we have also found some dominant CTL epitopes such as pp1a-38 and −84. In the process of finding dominant epitopes, we showed the existence of an immunodominance hierarchy in CD8^+^ T cell responses to these epitopes. The immunodominance hierarchy composed of multiple stages may offer us more variations in the epitope selection for designing CTL-based COVID-19 vaccines. These data may provide important information for further studies of T cell immunity in COVID-19 and the development of preventive and/or therapeutic CTL-based vaccines against SARS-CoV-2.

## Materials and Methods

### Prediction of CTL epitopes

Four computer-based programs including SYFPEITHI (31), nHLAPred (32), ProPred-I (33), and IEDB (34) were used to predict HLA-A*02:01-restricted CTL epitopes derived from pp1a of SARS-CoV-2 (GenBank accession numbers: LC528232.1 & LC528233.1). As shown in Table 1, 82 peptides with superior scores were selected and were synthesized by Eurofins Genomics (Tokyo, Japan). Two control peptides, FMP_58-66_ (sequence: GILFGVFTL) (35) and HIV-pol_476-484_ (sequence: ILKEPVHGV) (36), were synthesized as well.

### Mice

We used HLA-A*02:01 transgenic mice (37) that express a transgenic HLA-A*02:01 monochain, designated as HHD, in which human β2-microglobulin is covalently linked to a chimeric heavy chain composed of HLA-A*02:01 (α1 and α2 domains) and H-2D^b^ (α3, transmembrane, and cytoplasmic domains). Six-to ten-week-old mice were used for all experiments. Mice were housed in appropriate animal care facilities at Saitama Medical University, and were handled according to the international guideline for experiments with animals. This study was approved by the Animal Research Committee of Saitama Medical University.

### Cell lines

RMA-S-HHD is a TAP2-dificient mouse lymphoma cell line, RMA-S (H-2^b^) transfected with the HHD gene (37). RMA-S-HHD was cultured in RPMI-1640 medium (Nacalai Tesque Inc., Kyoto, Japan) with 10% FCS (Biowest, Nuaille, France) and 500 μg/ml G418 (Nacalai Tesque Inc.)

### Peptide binding assay

Binding of peptides to the HLA-A*02:01 molecule was measured using RMA-S-HHD cells, as described (46). Briefly, RMA-S-HHD cells were pre-cultured overnight at 26°C in a CO_2_ incubator, and then pulsed with each peptide at various concentrations ranging from 0.01 μM to 100 μM for 1 hour at 26°C. After 3 hours’ incubation at 37°C, peptide-pulsed cells were stained with anti-HLA-A2 monoclonal antibody, BB7.2 (47), followed by FITC-labeled goat anti-mouse IgG antibody (Sigma-Aldrich, St. Louis, MO). Mean fluorescence intensity (MFI) of HLA-A2 expression on the surface of RMA-S-HHD cells was measured by flow cytometry (FACSCanto™ II, BD Biosciences, Franklin Lakes, NJ), and standardized as the percent cell surface expression by the following formula: % relative binding = [{(MFI of cells pulsed with each peptide) – (MFI of cells incubated at 37°C without a peptide)}/{(MFI of cells incubated at 26°C without a peptide) – (MFI of cells incubated at 37°C without a peptide)}] × 100. The concentration of each peptide that yields the 50% relative binding was calculated as the half-maximal binding level (BL_50_). FMP_58-66_ and HIV-pol_476-484_ were used as control peptides.

### Peptide-encapsulated liposomes

Peptide-encapsulated liposomes were prepared using Lipocapsulater FD-U PL (Hygieia BioScience, Osaka, Japan) according to the manufacturer’s instructions with a slight modification. In brief, each of 54 synthetic peptides was dissolved in DMSO at a final concentration of 10 mM. For the first screening, the same amounts of 4 to 5 kinds of 10 mM peptides were mixed to make a total 100 μl, which was then diluted with 1.9 ml of H_2_O. For the second screening, 20 μl of each peptide at 10 mM was diluted to 2 ml with H2O. For the identification of dominant epitopes among the 10 peptides selected, 20 μl of each of the 10 peptide solutions at a concentration of 10 mM was mixed together, and the total volume was increased to 2 ml by adding 1.8 ml of H_2_O. The peptide solution was added into a vial of Lipocapsulater containing 10 mg of dried liposomes, and incubated for 15 min at room temperature. The resultant solution contains peptide-encapsulated liposomes.

### Immunization

Mice were immunized s.c. once or twice at a one-week interval with peptide-encapsulated liposomes (100 μl/mouse) together with CpG-ODN (5002: 5’-TCCATGACGTTCTTGATGTT-3’, Hokkaido System Science, Sapporo, Japan) (5 μg/mouse) in the footpad.

### Intracellular cytokine staining (ICS)

ICS was performed as described (30). In brief, after 1 wk following immunization, spleen cells were incubated with 50 μM of each peptide for 5 hours at 37°C in the presence of brefeldin A (GolgiPlug™, BD Biosciences), and were stained with FITC-conjugated anti-mouse CD8 monoclonal antibody (mAb) (BioLegend, San Diego, CA). Cells were then fixed, permeabilized, and stained with phycoerythrin (PE)-conjugated rat anti-mouse IFN-γ mAb (BD Biosciences). After washing the cells, flow cytometric analyses were performed using flow cytometry (FACSCanto™ II, BD Biosciences).

### Detection of CD107a and CD69 molecules on CD8^+^ T cells

For the detection of CD107a, spleen cells of immunized mice were incubated with 50 μM of each peptide for 6 hours at 37°C in the presence of monensin (GolgiStop™, BD Biosciences) and 0.8 μg of FITC-conjugated anti-mouse CD107a mAb (BioLegend). Cells were then stained with PE-Cy5-conjugated anti-mouse CD8 mAb (BioLegend). For the examination of CD69 marker, spleen cells of immunized mice were stimulated with 50 μM of each peptide for 4 hours at 37°C. Cells were then stained with PE-conjugated anti-mouse CD69 mAb (BioLegend) and FITC-conjugated anti-mouse CD8 mAb. Stained cells were analyzed by flow cytometry (FACSCanto™ II, BD Biosciences).

### *In vivo* CTL assay

*In vivo* CTL assay was carried out as described (46). In brief, spleen cells from naive HHD mice were equally split into two populations. One population was pulsed with 50 μM of a relevant peptide and labeled with a high concentration (2.5 μM) of carboxyfluorescein diacetate succinimidyl ester (CFSE) (Molecular Probes, Eugene, OR). The other population was unpulsed and labeled with a lower concentration (0.25 μM) of CFSE. An equal number (1 × 10^7^) of cells from each population was mixed together and adoptively transferred i.v. into mice that had been immunized once with a liposomal peptide. Sixteen hours later, spleen cells were prepared and analyzed by flow cytometry. To calculate specific lysis, the following formula was used: % specific lysis = [1-{(number of CFSE^low^ cells in normal mice)/(number of CFSE^high^ cells in normal mice)}/{(number of CFSE^low^ cells in immunized mice)/(number of CFSE^high^ cells in immunized mice)}] x 100.

### Statistical analyses

One-way ANOVA followed by post-hoc tests was performed for statistical analyses among multiple groups using Graphpad Prism 5 software (GraphPad software, San Diego, CA). A value of *P* < 0.05 was considered statistically significant.

## ACKNOWLEDGMENTS

The authors are grateful to Professor François A. Lemonnier (Pasteur Institute, Paris, France) for providing HHD mice and RMA-S-HHD cells. This work was supported by a Grant-in-Aid for Scientific Research (C) (JSPS KAKENHI Grant Number: JP18K06631) to M. M., a Grant-in-Aid for Early-Career Scientists (JSPS KAKENHI Grant Number: JP18K15430) to A. T. from Japan Society for the Promotion of Science, and a Grant from Ochiai memorial award 2018 to A. T. The authors have no conflicting financial interests.

